# Animate-inanimate object categorization from minimal visual information in the human brain, human behavior, and deep neural networks

**DOI:** 10.1101/2025.06.02.657367

**Authors:** Céline Spriet, Farzad Rostami, Jean-Rémy Hochmann, Liuba Papeo

## Abstract

The distinction between animate and inanimate things is a main organizing principle of information in perception and cognition. Yet, animacy, as a visual property, has so far eluded operationalization. Which visual features are necessary and sufficient to discriminate between animate and inanimate objects? At which level of the visual hierarchy does the animate-inanimate distinction emerge? Here, we show that the categorical boundary between animate and inanimate objects is preserved even among images of objects that are made unrecognizable and only retain the low- and mid-level visual features of their natural version. In three experiments, healthy human adults saw rapid sequences of images (6 Hz) where every five exemplars of a category (e.g., animate), an exemplar of another category (i.e., inanimate) was shown (1.2 Hz). Using frequency-tagging electroencephalography (ftEEG), we found significant neural responses at 1.2 Hz, indicating rapid and automatic detection of the periodic categorical change. Moreover, such effect was found––although increasingly weaker––for ‘impoverished’ stimulus-sets that retained only certain (high-, mid- or low-level) features of the original colorful images (i.e., grayscale, *texform* and phase-scramble images), and even if the images were unrecognizable, as tested with an object recognition task. Similar effects were found with Deep Neural networks (DNNs) presented with the same stimulus-sets. Reliable categorization effects for dramatically impoverished and unrecognizable images, in humans’ EEG and DNN data, demonstrate that the animate-inanimate distinction emerges early in the visual hierarchy and is remarkably resilient to the loss of visual information.

## Introduction

An astonishing capacity for *life detection* underlies animals’ survival. Humans and other species are endowed with mechanisms for rapid and accurate discrimination between animate and inanimate *things*. Accordingly, categorization by animacy (or animate-inanimate categorization) is a main organizing principle for object representation in the human brain (Warrington and Shallice, 1984; Caramazza and Shelton, 1998; Martin, 2007; Grill-Spector and Weiner, 2014), the first to emerge in infancy (Ayzenberg & Behrmann, 2024; Pauen, 2002; Spriet et al., 2022), and one of the most efficient mechanisms of visual perception (New et al., 2007; Thorpe et al., 1996; VanRullen & Thorpe, 2001).

Categorization by animacy is largely supported by visual information extracted within milliseconds, even from static images (Carlson et al., 2013; Cichy et al., 2014; Contini et al., 2017; Peykarjou et al., 2024b; Proklova et al., 2019). This ability is thought to rely on tuning to complex properties represented in higher-level visual areas, including the presence of a face/body, being alive, having agency and/or being unpredictable, among others (Bracci & Op De Beeck, 2023; Connolly et al., 2012; Jozwik et al., 2022; Leys et al., 2025; Papeo et al., 2017; Ritchie et al., 2021; Sha et al., 2015; Thorat et al., 2019). The kinds and degree of these properties are thought to determine an object’s position along an *animacy continuum*, reflecting the internal organization of the animate representational space.

Another line of research shows that the neural animate–inanimate distinction can also be supported by more basic visual information, carried by mid-level features such as shape and texture (Ayzenberg & Behrmann, 2022; Dehn et al., 2025; Li & Bonner, 2020; Loke et al., 2025; Long et al., 2017; Wang et al., 2022; for converging behavioral evidence see e.g., Schmidt et al., 2017; Tiedemann et al., 2022; Zachariou et al., 2018), and low-level features represented in the power-spectrum (Coggan et al., 2016; Grootswagers et al., 2017; Peykarjou et al., 2024b; Rice et al., 2014).

The present study asks whether basic visual features are themselves sufficient to elicit the animate-inanimate discrimination. In particular, we systematically assessed the animate-inanimate categorization across multiple stimulus manipulations that selectively impacted (high-, mid- or low-level) visual features, we quantitatively compared the resulting responses, and related them to behavioral categorization performance and representations at different layers of deep neural networks (DNNs). This systematic approach allowed us to assess how each feature considered here contributes to the animate-inanimate distinction, both independently and relative to the other features, across human brain responses, human behavioral judgments, and DNN representations.

Neural categorization responses were measured using frequency-tagging electroencephalography (ftEEG). In this paradigm, images from one category (e.g., inanimate) were presented in rapid sequence at 6 Hz, eliciting steady-state visual evoked potentials at the same frequency (Norcia et al., 2015). Every fifth image (the ‘oddball’) belonged to the other category (animate), introducing a periodic categorical change at 1.2 Hz. If animates are represented as distinct from inanimates, this categorical change should elicit a distinct neural response at 1.2 Hz (Quek & Rossion, 2017; Rossion et al., 2015; Rostami et al., 2026). We examined this response across experiments and conditions using realistic images of real-world objects and manipulated images that retained low-level (spectral power, color, contrast, luminance, number of pixels) or mid-level features (global form, texture).

We further investigated how neural categorization responses relate to participants’ ability to distinguish animate from inanimate stimuli under explicit judgment, including for the impoverished stimulus sets, and to stimulus representation across successive layers of a deep neural network (DNN), also for the impoverished sets. This latter analysis provides an image-computable measure of the animacy-related information retained by each stimulus transformation. Moreover, because DNNs generally predict object representations in human visual cortex, but have also been reported to show discrepancies in animacy recognition (Bracci et al., 2019; Duyck et al., 2024), we could assess the extent to which representations converge—or diverge—between biological and artificial vision.

## Materials and methods

Across five experiments, we combined complementary approaches to test the resilience of the animate–inanimate distinction to the loss of visual information. In Experiments 1–3, we used ftEEG to measure neural responses time-locked to stimulus presentation, and tested whether they encode the animate–inanimate distinction for intact image sets, compared with impoverished sets preserving only a subset of visual features: phase-scramble stimuli (Experiment 1; retaining color, contrast, luminance, power spectrum, and pixel distribution), grayscale phase-scramble stimuli (Experiment 2; all of the above except color), and texforms (Experiment 3; retaining texture and global shape). In Experiment 4, we assessed the participants’ ability to distinguish animate from inanimate stimuli, including for the impoverished stimulus sets, when an explicit judgment was required. In Experiment 5, we examined whether the visual information preserved in the impoverished sets supports the representation of animate and inanimate objects across successive layers of the deep convolutional neural network VGG-19 (Simonyan & Zisserman, 2015). Supplementary analyses tested and generalized the effects to different architectures (ResNet-50; He et al., 2016; Alexnet; Krizhevsky et al., 2012; CORnet-S; Kubilius et al., 2019; VGG16; Simonyan & Zisserman, 2015).

### Experiments 1-3

#### Participants

Experiments 1-3 involved a total of 36 healthy adults, with normal or corrected-to-normal vision and no history of psychiatric or neurological conditions. In particular, Experiment 1 involved 12 healthy adults (7 identified their gender as female, 5 as male, mean age 25.7 ± 4.7 years). One additional participant was tested and excluded from the analysis for falling asleep during the experiment. Experiment 1 was exploratory with respect to the sample size, and results were used to select the sample size for the following experiments. A sample size of 12 was selected taking into account a power analysis estimating that a sample size of 6 would be sufficient to observed a difference between intact *vs.* phase-scramble images comparable to Peykarjou et al. (2024b) (effect size Cohen’s d = 2.17, ß= 0.95, α = 0.05; GPower 3.1).

A power analysis based on Experiment 1 estimated that a sample size of 6 was required to obtain the difference between intact *vs.* phase-scramble images observed in Experiment 1 (phase-scramble condition; Cohen’s *d* = 2.05, ß= 0.95, *α* = 0.05; GPower 3.1). Therefore, in Experiments 2, we kept a sample of 12 as in Experiment 1 (9 identified their gender as female, 3 as male, mean age 26.3 ±6.3 years). One additional participant was tested and excluded from the analysis for falling asleep during the experiment. Likewise, a sample of 12 healthy adults was included in Experiment 3 (9 identified their gender as female, 3 as male, mean age 23.7 ± 2.7 years). One additional participant was tested and excluded from the analysis for falling asleep during the experiment.

Two independent samples of 19 native-French speakers (for Grayscale phase-scramble stimuli: *n* =19; 17 identified their gender as female, 2 as male, mean age 24.0 ± 3.8 years; for Texform stimuli*: n* =19; 17 identified their gender as female, mean age 23.3 ± 3.8 years) took part in an online behavioral-judgement task for assessing stimulus recognizability (see *Experiment 2: Impoverished set* below). All reported to have normal or corrected-to-normal vision and no history of psychiatric or neurological conditions.

All experiments were approved by the local ethics committee (CPP Ile de France VIII).

#### Stimuli

Each ftEEG experiment involved two conditions, one with an ‘intact’ stimulus set and the other with an ‘impoverished’ stimulus set. In Experiment 1, the ‘intact’ set consisted of 640 colorful images (hereafter, *original* stimuli), and the corresponding ‘impoverished’ set consisted of phase-scramble versions of the original set (hereafter, *phase-scramble* stimuli). In Experiment 2, the intact’ and ‘impoverished’ sets consisted of 640 grayscale images of animate and inanimate objects (hereafter, *grayscale* stimuli) and the corresponding phase-scramble versions (hereafter, *grayscale phase-scramble* stimuli). In Experiments 3, the ‘intact’ and ‘impoverished’ sets involved 350 grayscale images (hereafter, *grayscale* stimuli and the corresponding *texforms*, respectively).

##### Experiment 1: Intact set

The original set included 640 colorful photographs of real-world animate (*n*=320) and inanimate (*n*=320) objects taken from the internet, to broadly sample the variety of objects in the real world. The animate set included 51 fish, 70 birds, 179 nonhuman mammals, 14 amphibians and 6 turtles, all different from one another. Humans were excluded to prevent a bias in the animate-inanimate categorization. Insects, spiders and reptiles (except for turtles) were excluded to prevent emotional reactions (e.g., disgust or fear). The inanimate set included 223 artificial objects (16 exemplars of buildings and constructions, 108 exemplars of clothes, pieces of jewelry, buttons, coins, and tools, 66 pieces of furniture and 33 vehicles) and 97 natural objects (54 different fruits and vegetables, and 43 different flowers, bushes and trees) of big (41%; e.g., planes, trees) or small (59%; e.g., cups, fruits) real-world size. Objects (animate and inanimate) could appear in all sorts of orientations (1% of animals appeared in front view, showing a full face), and were all resized to fit a box of 629×629 pixels with a gray background (Figure 1A), which subtended a visual angle of 16° from a distance of ∼60 cm. Relatively large image size was used to make sure that stimuli were visible even under fast and low contrast stimulus presentation (de Heering & Rossion, 2015).

**Figure 1.**
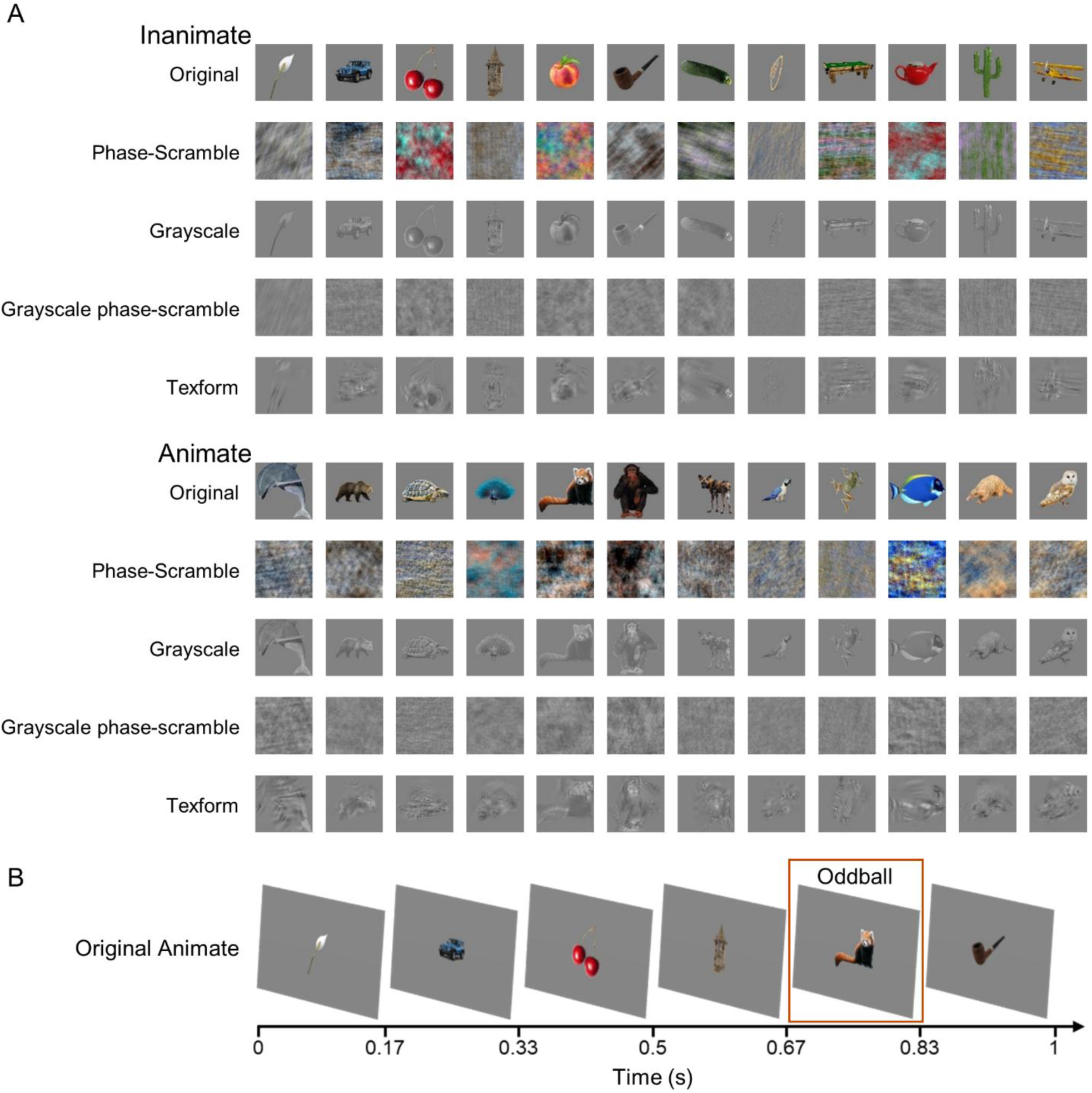
Illustration of stimuli and experimental design in Experiments 1-3. (A) Examples of stimuli from the *original set* (Experiment 1) in the inanimate condition (top) and animate condition (bottom) and the different versions of those images: phase-scramble (Experiment 1), grayscale (Experiments 2-3), grayscale phase-scamble (Experiment 2) and texforms (Experiment 3). (B) One-second extract from the “animate” stimulation sequence testing the categorization of animates (*oddball* category) among inanimate objects (*standard* category), with the *original set*.

Two types of stimulation sequences were created, ‘animate’ and ‘inanimate’, named after the category that served as *oddball*, i.e., the less frequent stimulus-category used to elicit the periodic categorization-response (Figure 1B). Thus, in the animate-sequence, 320 inanimate objects were shown as *standard* (i.e., the frequent stimulus-category in the sequence), and 68 animate objects (9 fish, 13 birds, 39 nonhuman mammals, 4 amphibians and 3 turtles) were shown as *oddball*; in the inanimate-sequence, 320 animate objects were shown as standard, and 68 inanimate objects (4 buildings or constructions, 19 clothes/jewels/buttons/coins/tools, 10 exemplars of furniture and 2 vehicles, 20 fruits/ vegetables, and 13 flowers/bushes/trees) were shown as *oddball*. Within each sequence, standard- and oddball-stimuli were matched for number of informative pixels (i.e., number of colorful pixels that defined the actual object, excluding the gray pixels that belong to the background; animate-sequence: 95% CI = -2.490 – 0.841; *t*(386) = - 0.973; *P* = .331; inanimate-sequence: 95% CI = -3.137 – 0.635; *t*(386) = -1.305; *P* = .193).

##### Experiment 1: Impoverished set

Phase-scramble stimuli were created by manipulating the spatial spectrum of the original stimuli through phase-scrambling, using the *imscramble* function (http://martin-hebart.de/webpages/code/stimuli.html) in Matlab (The Mathworks, Natick, MA). This manipulation preserved low-level information including the number of pixels, color, contrast, luminance and power-spectral distribution of the original images, but removed mid-level (shape and texture) information, effectively rendering the object identity unrecognizable. Stimulation sequences were created as for the intact set.

##### Experiment 2: Intact set

Grayscale images were obtained from the ‘intact’ set (original stimuli) of Experiment 1, using the step1_lumContrastOriginals_fromGreen function (https://github.com/brialorelle/texformgen). Stimulation sequences were created as in Experiment 1.

##### Experiment 2: Impoverished set

From the intact set of Experiment 2, phase-scramble stimuli were created as in Experiment 1, to remove mid-level (shape and texture) information, while preserving the number of pixels, contrast, luminance and power-spectral distribution of the grayscale images. We verified that the objects in these images were unrecognizable with a naming task administered to an independent sample of 19 native-French speakers participants, recruited and tested on the online platform Testable.org (Rezlescu et al., 2020). An image was considered ‘unrecognizable’ when no more than one participant named it correctly at the basic-category level (e.g., ‘fish’, ‘flower’, etc.). All images resulted to be ‘unrecognizable’ (only two images were named correctly by one participant; all others were never correctly named).

##### Experiment 3: Impoverished set

Texforms are modified versions of the grayscale images which preserve partial information about texture and form (Freeman & Simoncelli, 2011; Long et al., 2017). Following the method in Deza et al. (2019), 640 texforms were created from the 640 grayscale stimuli and their recognizability was assessed with the online behavioral-judgement task. Based on this study, for Experiment 3, we selected a subset of 195 animate and 175 inanimate unrecognizable texforms (texforms that were correctly at the basic-category level by no more than one participant; Long et al., 2017). This way, we could test categorization based on the visual features preserved in the texforms, without the images being recognized at a higher level. To match the number of items between conditions, we randomly removed 20 animate images, resulting in a total of 350 stimuli (175 animates and 175 inanimates). In the animate-sequence, 175 inanimate objects were shown as *standard*, and 47 animate objects were shown as *oddball*; in the inanimate-sequence, 175 animate objects were shown as standard, and 47 inanimate objects were shown as *oddball*.

##### Experiment 3: Intact set

From the ‘intact’ set of Experiment 2 (grayscale stimuli), we picked the 175 inanimate and 175 animate stimuli corresponding to the texforms selected above. Sequences were created as for the impoverished set.

#### Procedures

##### Paradigm

In Experiments 1-3, participants sat on a chair ∼60 cm away from a 60 Hz computer screen (resolution 1920×1200 pixels, size 51.5×32.2cm), where stimuli were presented centrally (16° of visual angle). Each trial lasted 32 s, including a 2-s fade-in phase (increase in contrast) and a 2-s fade-out phase (decrease in contrast), to ease the stimulus presentation for the participant’s eyes, and a sequence of stimuli, presented at a base frequency of 6 Hz (6 images per *s*; 166.67 ms per image), in a squarewave design, where every 5 images of one category, one ‘oddball’ image of the other category was shown (Figure 1 B). Rapid serial visual presentation combined with EEG is used to elicit a steady-state visual evoked potential (SSVEP) at the stimulation (“base”) frequency, and a second response at the “oddball” frequency when oddball stimuli consistently recruit a distinct neuronal population. This frequency-tagging EEG (ftEEG) paradigm is particularly well suited to address our question on the animate-inanimate distinction as it directly captures the two defining aspects of categorization: within-category generalization and between-category discrimination. Specifically, an oddball response can only emerge when oddball stimuli elicit a common neural response that differs from the response elicited by the standard (base frequency) stimuli. Thus, the presence of an oddball response provides evidence that the neural system both generalizes across exemplars within a category and discriminates that category from the standard category. Importantly, ftEEG provides a high signal-to-noise ratio (SNR) because the neural responses of interest are isolated at predefined stimulation frequencies (here, 6 and 1.2 Hz). This sensitivity is especially valuable when investigating potentially subtle category-selective responses under visually noisy conditions (e.g., visually degraded stimuli). Moreover, because stimuli are presented rapidly under strong forward- and backward-masking, ftEEG is particularly well suited to probing automatic categorization processes (de Heering & Rossion, 2015; Norcia et al., 2015; Peykarjou, 2022; Retter et al., 2020; Rossion & Boremanse, 2011).

In our sequences, the five standard stimuli that preceded each oddball were selected pseudo-randomly to prevent repetitions of the same image in a trial and the presentation of two images of the same subordinate-level category at the same frequency as the oddball presentation (e.g., a dolphin and, after five images, another mammal, e.g., a cat). At the end of each trial, a test-image from the standard category was shown and participants had to report whether it was shown during the trial, by pressing “A” or “P” (“yes” or “no”). This task was included to motivate participants to pay attention to the stimuli.

Each participant was tested in two conditions, involving an ‘intact’ (i.e., original or grayscale stimuli) and an ‘impoverished’ (i.e., phase-scramble, grayscale phase-scramble or texform stimuli) set, respectively. Each condition included 16 trials of animate-sequences (i.e., animate-*oddballs* among inanimate-*standards*) and 16 trials of inanimate-sequences (i.e., inanimate-*oddballs* among animate-*standards*) for a total of 64 trials (16 trials x 2 conditions x 2 types of sequences). For each participant, the same list of images was used in the two conditions.

The 16 trials of a condition were presented in a single block, yielding a total of four blocks (animate and inanimate sequences in intact and impoverished conditions). The two blocks of the same condition were presented one after the other, with the order of type of sequence (animate or inanimate) and condition (intact or impoverished) counterbalanced across participants for Experiments 1 and 2. For Experiment 3, to prevent recognition based on previous exposure to ‘intact’ images, blocks with texform stimuli were always presented first. Stimulus presentation was controlled using Psychtoolbox (Brainard, 1997) through Matlab. The whole experiment lasted ∼45 minutes.

##### EEG recording

EEG data were acquired using 128-channel EGI nets (Electrical Geodesics, Inc.). Data were acquired with vertex reference, using the EGI Net Station acquisition software, continuously digitized at a sampling rate of 1kHz (net amp 400 system EGI). Impedance was lowered for each participant as much as possible, and did not exceed 40kΩ. During the experiment, triggers were sent from the experimental computer to the acquisition computer via a light sensor: a white square appeared in correspondence to the sensor (i.e., at the bottom right of the screen) at the beginning and at the end of each trial. This manipulation allowed a high precision in timing the beginning and the end of a trial, allowing to extract the EEG data corresponding to the presentation of the trial and time-locking EEG data to the stimuli presentation.

### EEG analysis

#### EEG preprocessing

Data preprocessing was performed with the EEGLAB toolbox (Delorme & Makeig, 2004) and MATLAB R2015b. Raw data for each participant were filtered using a 4th-order high pass butterworth filter at 0.1 Hz and a 4th-order low pass butterworth filter at 100 Hz. All electrodes were re-referenced using the average of all electrodes as a reference. Data were segmented by trials, taking 25 s for each trial (30 complete cycles) starting 2.5 s after the trial onset and excluding the fade-in and fade-out phases. Trials were averaged separately for each participant, each condition (intact and impoverished) and each type of sequence (animate and inanimate), in the time-domain.

#### EEG frequency domain analysis

Data were analyzed considering the two conditions (‘intact’ and ‘impoverished’, thereafter referred to as ‘Image Type’), averaging across types of sequences (animate, inanimate, thereafter referred to as ‘Sequence Type’). A Fast Fourier Transform (FFT) was applied for data examination in the EEG frequency-domain at the high frequency resolution of 0.04 Hz (1/25 seconds), inspecting the response at the oddball frequency and harmonic frequencies (i.e., multiple integers of the fundamental frequency of 1.2 Hz: 2.4, 3.6, 4.8, etc.), resulting from the FFT decomposition of the EEG signal.

In the first analysis, we examined whether, for each experiment, for each condition (intact and impoverished), the amplitude of the responses at the base and oddball frequencies were significantly higher than the noise level (i.e., significantly greater than 0; one-sample *t* tests, one tail). For this analysis, responses at the base or oddball frequency were determined independently for the two conditions, by considering the harmonics with a *z*-score higher than a threshold of 1.64 (*α* = .05, one-tailed; see Table 1 for the *z*-score of the oddball harmonics) and electrodes with a mean z-score higher than 3.33 (α = .0004, one-tailed, Bonferroni-corrected for 128 electrodes). Following established procedures (de Heering & Rossion, 2015; Rekow et al., 2022; Retter et al., 2020; Rossion et al., 2015), we computed group-level z-scores for all harmonics of the base frequency below 50 Hz and for all harmonics of the oddball frequency below 12 Hz, excluding the base frequency and its harmonics. Group-level frequency spectra were obtained separately for each condition (intact and impoverished) by averaging participants’ EEG signals in the time domain, applying the FFT, and then averaging across sequence types and electrodes. Z-scores were computed by subtracting the mean amplitude of the local baseline from the amplitude at each frequency of interest and dividing the result by the standard deviation of the local baseline. The local baseline was defined as the mean of amplitudes of the 24 surrounding frequency bins (12 on each side excluding the immediate adjacent frequency bin for a frequency range of ±0.48 Hz) excluding the maximum and minimum (Quek & Rossion, 2017; Retter & Rossion, 2016). With this procedure, a maximum of 8 harmonics were selected for each frequency in all experiments (see Table 1; the selected harmonics for each experiment and condition are in highlighted bold). Electrodes were selected by computing group-level z-scores at each electrode, averaged across the selected harmonics. For statistical analyses, baseline-subtracted amplitudes were computed for each participant, condition, and electrode, summed across selected harmonics and averaged across selected electrodes. Baseline-subtracted amplitudes were computed by subtracting the mean amplitude of the local baseline from the amplitude at each frequency of interest.

**Table 1.**
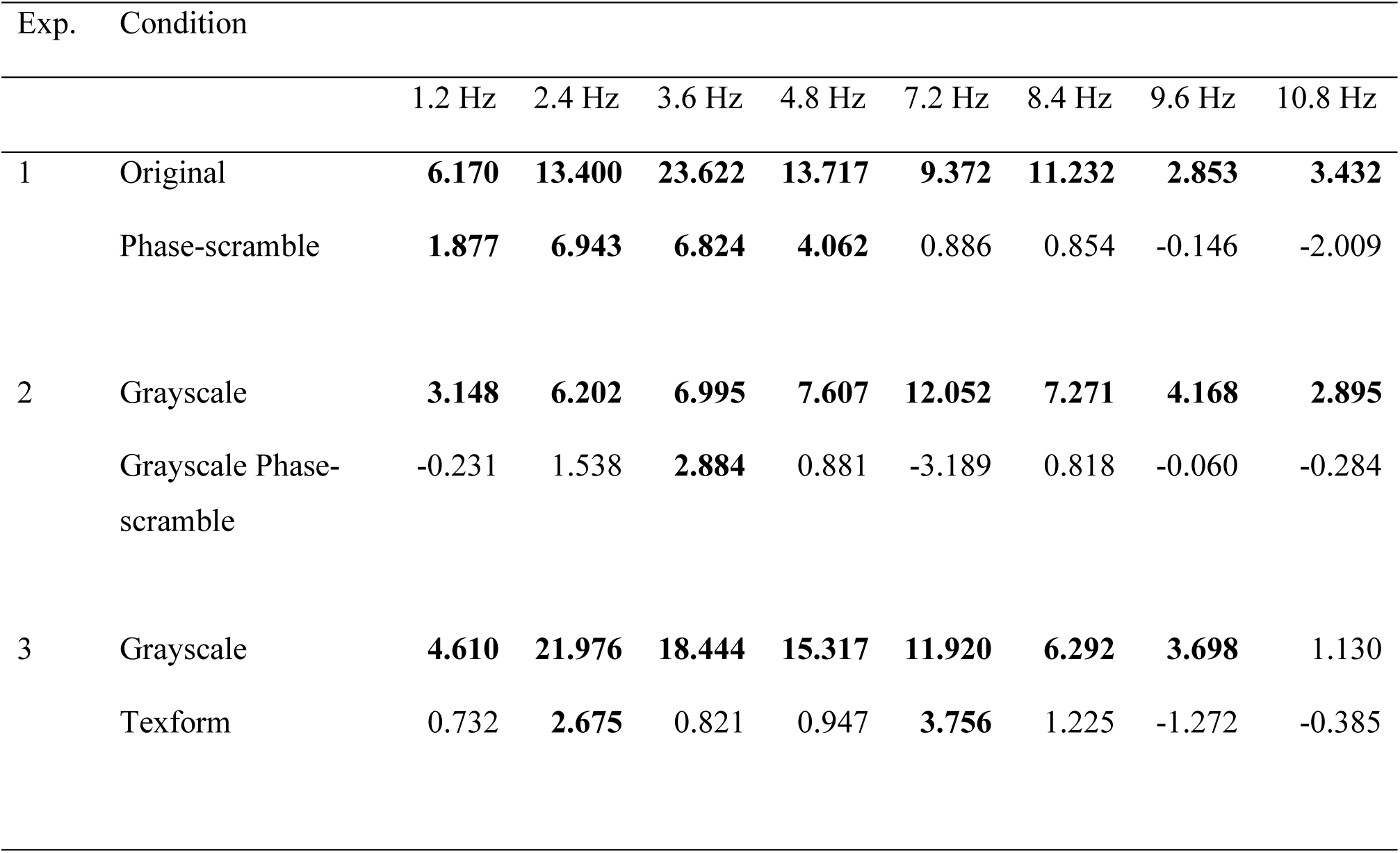
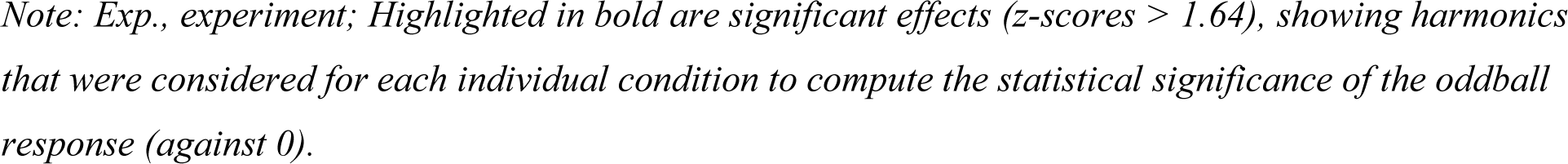
Categorization-response in Experiments 1-3. Z-scores of the first eight harmonics (excluding the 5^th^) reflecting the oddball frequency and for each condition of Experiments 1-3.

| Exp. | Condition | 1.2 Hz | 2.4 Hz | 3.6 Hz | 4.8 Hz | 7.2 Hz | 8.4 Hz | 9.6 Hz | 10.8 Hz |
| --- | --- | --- | --- | --- | --- | --- | --- | --- | --- |
| 1 | Original | **6.170** | **13.400** | **23.622** | **13.717** | **9.372** | **11.232** | **2.853** | **3.432** |
|  | Phase-scramble | **1.877** | **6.943** | **6.824** | **4.062** | 0.886 | 0.854 | -0.146 | -2.009 |
| 2 | Grayscale | **3.148** | **6.202** | **6.995** | **7.607** | **12.052** | **7.271** | **4.168** | **2.895** |
|  | Grayscale Phase-scramble | -0.231 | 1.538 | **2.884** | 0.881 | -3.189 | 0.818 | -0.060 | -0.284 |
| 3 | Grayscale | **4.610** | **21.976** | **18.444** | **15.317** | **11.920** | **6.292** | **3.698** | 1.130 |
|  | Texform | 0.732 | **2.675** | 0.821 | 0.947 | **3.756** | 1.225 | -1.272 | -0.385 |

Note: Exp., experiment; Highlighted in bold are significant effects (z-scores > 1.64), showing harmonics that were considered for each individual condition to compute the statistical significance of the oddball response (against 0).

Second, we examined whether the amplitude of the responses at the base and oddball frequencies differed between the intact and impoverished conditions of each experiment. To allow the comparison, responses for the two conditions of each experiment were extracted from same harmonics and electrodes. In particular, baseline-subtracted amplitudes were summed across all harmonics showing an effect in at least one of the two conditions (Image Type), and averaged across all 128 electrodes. Additionally, using the same data, a 2 Image Type x 2 Sequence Type repeated-measures ANOVA was used to test whether the oddball response of the Image Type (intact and impoverished) was affected by the Sequence Type (animate or inanimate).

Finally, we examined whether the amplitude of the responses at the base and oddball frequencies differed across experiments. Again, to allow the comparison, responses for all conditions of all experiments were extracted from same harmonics and electrodes. In particular, for each participant, for each experiment and each condition, the response was computed as the sum of baseline-subtracted amplitudes of all the 8 harmonics (1.2 Hz; 2.4 Hz; 3.6 Hz; 4.8 Hz; 7.2 Hz; 8.4 Hz; 9.6 Hz and 10.8 Hz), averaged across all 128 electrodes and across the sequence types (animate, inanimate). These values were entered in an ANOVA with Image Type (intact, impoverished) as within-subjects factor and Experiment (1, 2 3) as between-subjects factor.

### Experiment 4

#### Participants

Experiment 4 involved a total of 60 healthy, adult, native-English speakers, external to the above EEG and naming studies. They were recruited online through the platform Testable.org (Rezlescu et al., 2020), and randomly assigned to one of three tasks involving, respectively, the phase-scramble stimuli of Experiment 1 (*n*=20, 12 identified their gender as female, 8 as male, mean age 26.6 ± 4 years), the grayscale phase-scramble stimuli of Experiment 2 (*n*=20, 7 identified their gender as female, 12 as male, 1 as other, mean age 26.5 ± 4.1) and the texforms used in Experiment 3 (*n*=20, 9 identified their gender as female, 11 as male, mean age 26.7 ± 3.3).

#### Experimental design

Participants were instructed to sit 60 cm away from the screen, align their eyes to the center of the screen, not move during the experiment, and turn off their phone and other devices. Prior to the experiment, they gave informed consent and followed instructions for screen calibration. They were instructed to performed a yes-no animal detection task (i.e., to decide whether each image depicted an animal or not) on 640 transformed images of existing objects. Stimuli were presented one by one in the center of the screen (16° of visual angle) and remained until the participant responded, by clicking on the “yes” or “no” button below the image, using the mouse. The side of the “yes” and “no” buttons were counterbalanced across participants. Images were presented in random order.

#### Statistical analyses

The yes-no animal detection task was used to test whether, even if objects in the impoverished sets were not recognizable at the basic-category level, as per the stimulus selection, participants could still *guess* the superordinate category (i.e., animate or inanimate). To this end, *d’* for each condition were computed as a measure of discrimination and tested against chance (one-sample *t*-test, one-sided), and compared across stimulus sets (two-sample *t*-test; α = .017, Bonferroni-corrected for 3 comparisons).

### Experiment 5

#### Stimuli and experimental design

We selected the convolutional deep neural network (DNN) VGG-19 (Simonyan & Zisserman, 2015), a computational model of hierarchical object processing in the primate ventral visual stream that achieves human-level object recognition performance, particularly in its deeper layers (Khaligh-Razavi & Kriegeskorte, 2014; Kheradpisheh et al., 2016; Kubilius et al., 2016; Yamins et al., 2014). We then examined the extent to which the impoverished stimulus sets preserved object representations—and, in particular, the animate–inanimate categorical structure—across successive layers of the network, relative to the representation of the original stimuli in the final fully connected layer. Specifically, for each stimulus set, we identified the DNN layer whose representations best matched those of the original stimulus set, thereby estimating whether the preserved information corresponded primarily to low-, mid-, or high-level stages of visual processing.

Multiple processing stages in this DNN transform input images through a series of nonlinear operations. Initially, convolutional layers apply kernels to small regions of the input image, capturing spatial hierarchies of visual features. Then, a rectified linear unit (ReLU) function introduces nonlinearity by thresholding activations at zero, allowing only positive activations to pass forward. Then, max pooling layers perform downsampling, which reduces the spatial dimensions of the input while preserving important features. Finally, fully connected layers flatten the processed input into a one-dimensional vector, which represents the class scores. This network was trained on approximately 1.2 million images from the ImageNet database (ILSVRC2012), encompassing 1,000 classes of objects including animals (40%) and inanimate objects (60%). We used a pretrained version of this model available in MATLAB (MatConvNet; Vedaldi & Lenc, 2015), with standardized preprocessing steps such as mean subtraction of the training images and scaling of all stimuli to 224×224 pixels.

#### Statistical analyses

We extracted features from convolutional layers (‘conv1_1’ through ‘conv5_4’) and fully connected layers (‘fc6’, ‘fc7’, and ‘fc8’). For each of these layers, for each set of stimuli used in Experiments 1-3, we computed a dissimilarity matrix (Kriegeskorte et al., 2008; see Figure 4 for examples) representing pairwise dissimilarities between the features extracted from the stimuli (1 minus *r*, where *r* is the *Pearson* coefficient of the correlation between vectors of features representing two stimuli). Separately for each stimulus set, we performed *Spearman* correlations between the matrix extracted from each layer and the *reference matrix*, corresponding to the dissimilarity matrix modeling the representation of stimuli in the original set, in the output layer of the DNN (‘fc8’). Significance was assessed with 1000 row/column permutations of the reference matrix. The resulting *p* values were corrected for multiple comparisons across the layer × stimulus-set RSA correlations using the Benjamini–Hochberg false discovery rate procedure. The reference matrix was chosen as it reflects the performance on the stimulus set with the highest complexity (i.e., highest number of features), and the closest to the stimuli used for training the DNN, thus providing the most reliable representation of the stimuli by the DNN, typically yielding the best performance of the model. While the reference matrix represented graded similarity relations between pairs of stimuli, we further verified that it also carried a binary animate-inanimate categorical model (see Supplementary Materials). The performance throughout all the layers of the DNN was also compared between the different stimulus sets, using pairwise *t*-tests on the Fisher-transformed correlation coefficients. The same analyses were repeated within different hierarchical architectures using the THINGvision package in python (Muttenthaler & Hebart, 2021; see Supplementary Materials).

## Results

### Experiment 1

Experiment 1 tested whether the fast presentation of a sequence of images of objects from a large and heterogeneous set elicited a response (i.e., baseline-corrected amplitude significantly greater than noise-level) in correspondence to the regular appearance of an “oddball” animate object among inanimate objects, and vice versa, thus signaling detection of the categorical change. By using phase-scramble versions of the same images, we tested to what extent such categorization-response could be also elicited by stimuli that only preserved low-level visual features of the objects, such as power-spectrum, color, contrast, and luminance.

First, we verified that participants showed a significant base response (6 Hz), i.e., a positive (higher than 0) amplitude at the base frequency, indicating reliable synchronization of neural activity with visual stimulation (‘intact’: *M_Amplitude_*±*sd* = 0.80±0.36; 95% CI = 0.62 – Inf; *t*(11) = 7.71; *P* < .001; *d* = 2.23; ‘impoverished’: *M_Amplitude_*±*sd* = 0.77±0.35; 95% CI = 0.59 – Inf; *t*(11) = 7.67; *P* < .001; *d* = 2.21). Then, we analyzed the response at the oddball frequency which reflects the automatic detection of the categorical change upon stimulus presentation. As described in the Methods, in Experiment 1 (as well as in Experiments 2-3), oddball responses for each condition were computed by averaging across the harmonics and electrodes that exceeded the significance threshold for that condition, as determined from the group-level data (Table 1). A significant (above noise-level) widespread categorization response, peaking over posterior electrodes, was found in both the ‘intact’ condition (*M_Amplitude_*±*sd* = 0.32±0.08; 95% CI = 0.28 – Inf; *t*(11) = 13.25; *P* < .001; *d* = 3.82; Figure 2 A) and the ‘impoverished’ condition (*M_Amplitude_*±*sd* = 0.13±0.06; 95% CI = 0.10 – Inf; *t*(11) = 7.48; *P* < .001; *d* = 2.16; Figure 2 A), but it was larger in the former condition (*M_Difference_*± *sd* = 0.21 ±0.10; 95% CI = 0.14 – 0.27; *t*(11) = 7.11; *P* < .001; *d* = 2.05). There was no difference between the two Sequence-Type conditions (animate-as-oddball *vs*. inanimate-as-oddball). Indeed, a 2 Image Type x 2 Sequence-Type ANOVA showed an effect of Image Type, *F*(1,11) = 51.77, *P* < .001, *η²* = .83, but no effect of Type of sequence, *F*(1,11) = 1.11, *P* = .32, *η²* = .09, or interaction, *F*(1,11) = 0.97, *P* = .35, *η²* = .08. As described in the Methods, in Experiment 1 (as well as in Experiments 2-3), oddball responses for comparisons between conditions were computed by averaging across the same harmonics and electrodes for all conditions.

**Figure 2.**
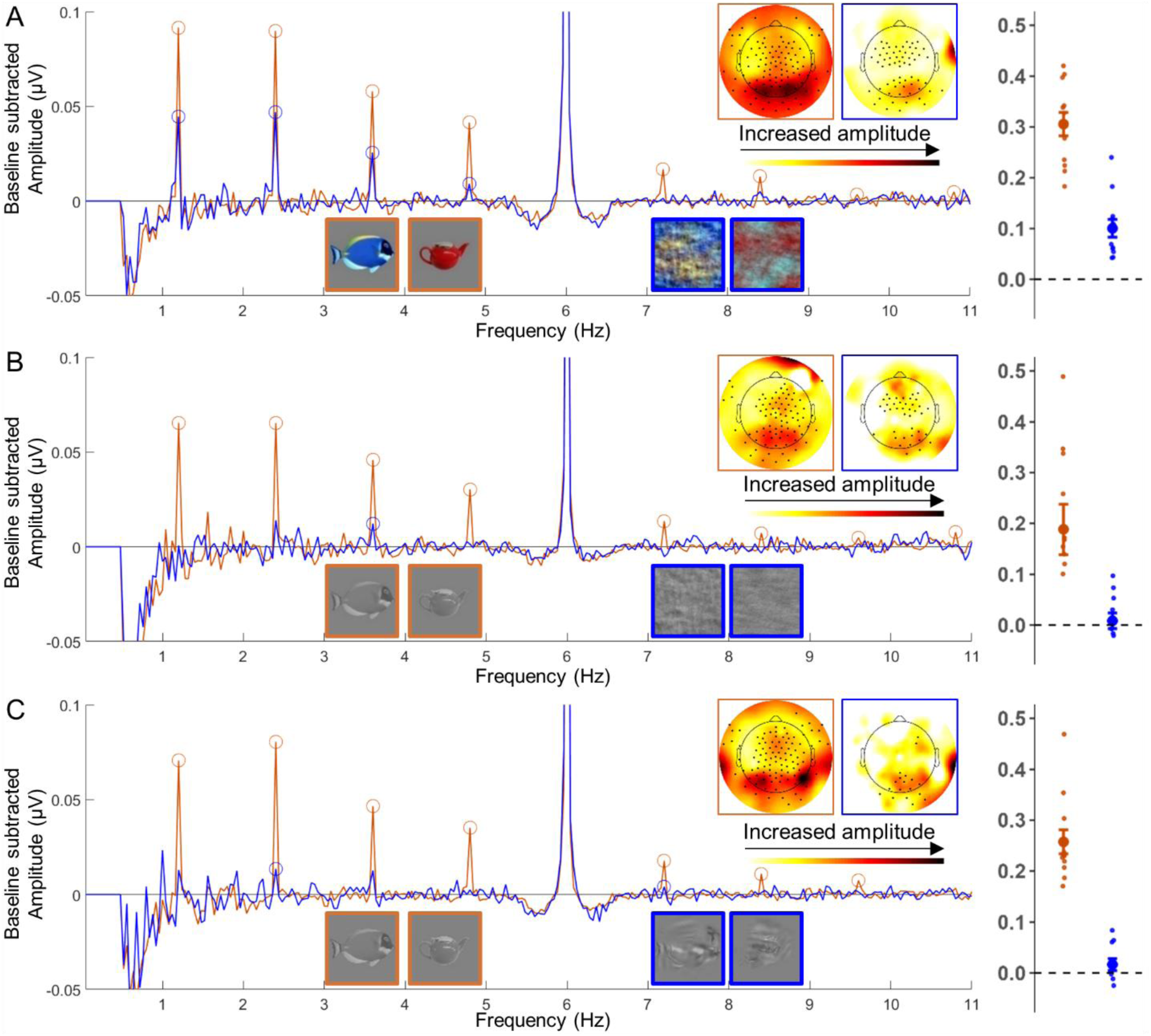
Responses at the oddball frequencies. in Experiments 1-3. *Left.* Baseline-subtracted amplitudes as a function of frequencies in the conditions with original (orange) and phase-scramble stimuli (blue) of Experiment 1 **(A),** with grayscale (orange) and grayscale phase-scramble stimuli (blue) of Experiment 2 (**B**), and with grayscale stimuli (orange) and texforms (blue) of Experiment 3 (**C**). Circles signal harmonics showing a significant oddball response for a condition. The scalp distribution of the group baseline-subtracted amplitudes summed over the significant harmonics (in circles) show response peaks over posterior electrodes. *Right*: Baseline-subtracted amplitudes summed over significant harmonics (highlighted by circles in the plots on the left) and averaged over significant electrodes (highlighted by black dots in the scalp distribution plot) for the ‘intact’ (orange) and ‘impoverished’ (blue) conditions in each experiment. Small dots denote individual responses, thicker dots denote group-average responses, lines denote standard errors of the mean.

#### Experiment 2

Results of Experiment 2 replicated those of Experiment 1, with grayscale (‘intact’) and grayscale phase-scramble (‘impoverished’) images, matched for contrast and luminance between the animate and inanimate set. We found a significant response at the base frequency in both conditions (grayscale: *M_Amplitude_* ±*sd* = 0.44 ±0.17; 95% CI = 0.35 – Inf; *t*(11) = 8.93; *P* < .001; *d* = 2.58; grayscale phase-scramble: *M_Amplitude_*±*sd* = 0.36 ±0.16; 95% CI = 0.28 – Inf; *t*(11) = 7.98; *P* < .001; *d* = 2.30). Importantly, a significant categorization response peaking over posterior electrodes was found in both the grayscale ‘intact’ condition (*M_Amplitude_* ±*sd* = 0.24±0.15; 95% CI = 0.16 – Inf; *t*(11) = 5.42; *P* < .001; *d* = 1.57; Table 1; Figure 2 B) and the ‘impoverished’ phase-scramble condition (*M_Amplitude_*±*sd* = 0.01 ±0.02; 95% CI = 0.00 – Inf; *t*(11) = 2.46; *P* = .016; *d* = 0.71; Figure 2 B; Table 1). The categorization response was larger in the ‘intact’ condition (*M_Difference_* ±*sd* = 0.18±0.17; 95% CI = 0.07 – 0.29; *t*(11) = 3.64; *P* = .004; *d* = 1.05), with no differences between Sequence Types, as shown by a 2 Image Type x 2 Sequence Type ANOVA (Effect of Image Type: *F*(1,11) = 12.87, *P* = .004, *η²* = .54; Effect of Sequence Type: *F*(1,11) = 0.01, *P* = .91, *η²* = .00; Interaction: *F*(1,11) = 0.20, *P* = .66, *η²* = .02).

#### Experiment 3

For both the grayscale stimuli (same as in Experiment 2) and the texforms, which only retained mid-level features of the objects (texture and global form), we found a response at the base frequency (grayscale: *M_Amplitude_* ±*sd* = 0.61±0.31; 95% CI = 0.45 – Inf; *t*(11) = 6.77; *P* < .001; *d* = 1.95; texforms: *M_Amplitude_* ±*sd* = 0.45±0.19; 95% CI = 0.36 – Inf; *t*(11) = 8.27; *P* < .001; *d* = 2.39) (Figure 2 C). Moreover, a categorization response (i.e., at the oddball frequency), peaking over posterior electrodes, was found for both grayscale stimuli (*M_Amplitude_* ±*sd* = 0.27±0.09; 95% CI = 0.22 – Inf; *t*(11) = 10.79; *P* < .001; *d* = 3.12) and texforms (*M_Amplitude_* ±*sd* = 0.02±0.02; 95% CI = 0.01 – Inf; *t*(11) = 2.67; *P* = .011; *d* = 0.77) (Figure 2 C), but it was larger in the former ‘intact’ condition (*M_Difference_* ±*sd* = 0.24±0.10; 95% CI = 0.17 – 0.30; *t*(11) = 8.21; *P* < .001; *d* = 2.37). There were no differences between Sequence Types as shown by a 2 Image Type x 2 Sequence Type ANOVA (Effect of Image Type: *F*(1,11) = 66.60, *P* < .001, *η²* = .86; Effect of Sequence Type: *F*(1,11) = 0.04, *P* = .85, *η²* = .00; Interaction: *F*(1,11) = 3.89, *P* = .074, *η²* = .26).

In sum, results showed a categorization response at the oddball frequency for all stimulus sets (Figure 2). This response was larger for the feature-richer (‘intact’) images than for the ‘impoverished’ ones. Because the impoverished images were more difficult to process, we considered whether participants showed reduced attention or engagement in these conditions related to the weaker categorization responses. Several observations speak against this possibility. First, participants’ performance on the post-sequence task (judging whether an image had appeared in the preceding sequence) was at chance in all conditions except the intact condition of Experiment 1 (see Supplementary Materials). However, this was not the only condition to produce a significant categorization response, or a response larger than its corresponding impoverished condition, indicating that performance on the post-sequence task—a proxy for attention to the stimuli—did not reliably predict the oddball response. Second, we examined the amplitude of the base-frequency response as another potential index of attention to the visual stimulation. This response did not differ between intact and impoverished conditions in Experiments 1 and 2 and differed only in Experiment 3 (intact grayscale stimuli > texforms; see Supplementary Materials). Yet, categorization responses were reduced for the impoverished stimuli in all three experiments. Thus, the base-frequency response likewise failed to account for the differences in categorization responses. Taken together, these findings suggest that any differences in attention or engagement between intact and impoverished conditions cannot explain the observed pattern of oddball-frequency responses.

### Comparison between Experiments 1-3

We compared the categorization-response across experiments, to investigate how it varied as a function of stimulus complexity. For each experiment, for each participant and condition, we computed a single response corresponding to the sum of baseline-subtracted amplitudes of the eight harmonics showing a significant categorization-response in at least one condition of one experiment (see Table 1), averaged across all 128 electrodes and across sequence types (animate-as-oddball, inanimate-as-oddball). Thus, for comparison across conditions and experiments, the same harmonics and electrodes were considered to compute the oddball responses. An ANOVA with Image Type (‘intact’ *vs.* ‘impoverished’) as within-subjects factor and Experiment as between-subjects factor showed effects of Image Type, *F*(1,33) = 94.77, *P* < .001, *η²* = .74, and Experiment, *F*(2,33) = 7.59, *P* = .002, *η²* = .32, but no interaction between the two, *F*(2,33) = 0.70, *P* = .51, *η²* = .04 (see Figure 3 A for a visualization of the response across Image Type and Experiment).

**Figure 3.**
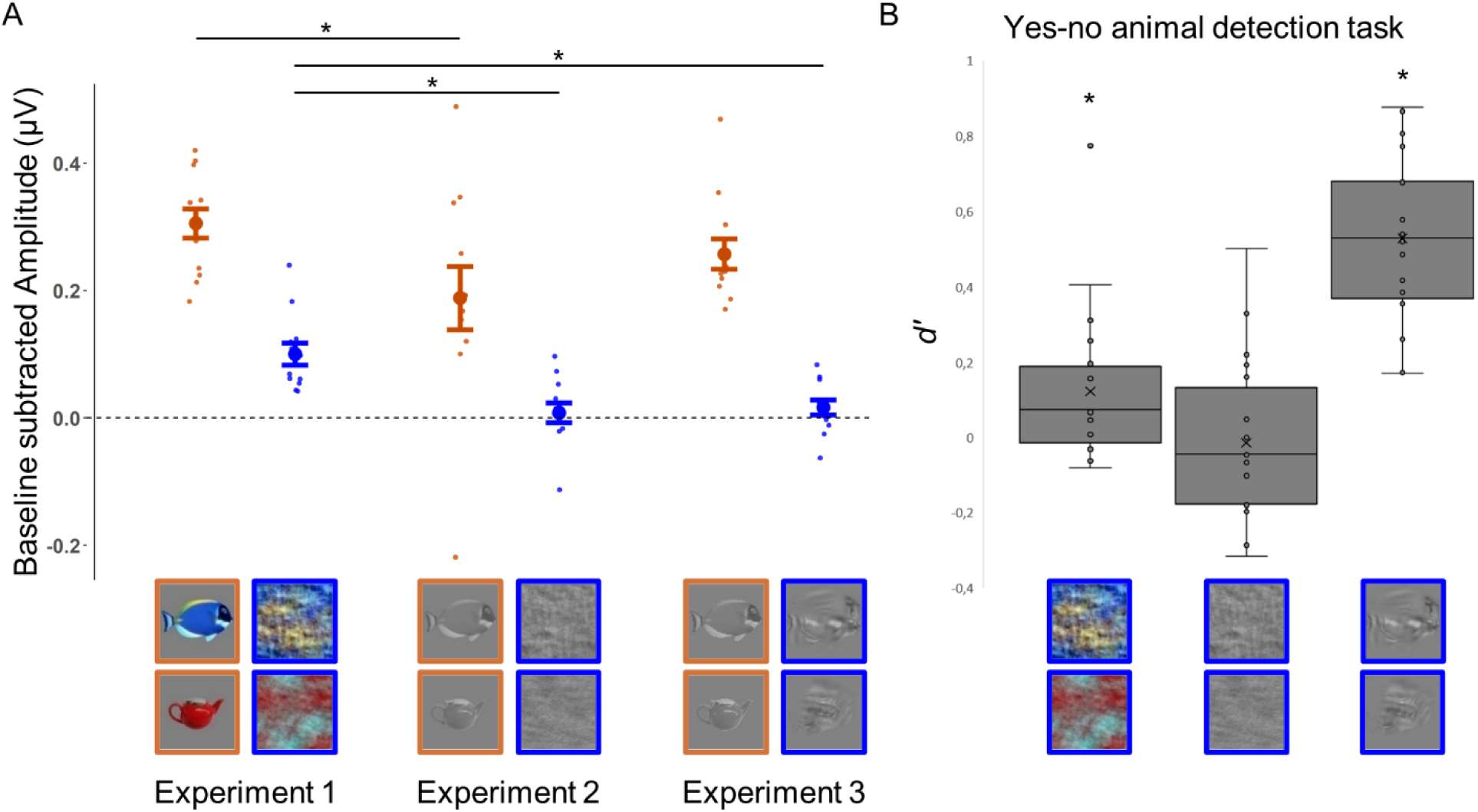
Comparison between Experiments 1-3. **(A)** Mean categorization response (sum of baseline subtracted amplitude of the same eight harmonics, averaged over all 128 electrodes, and standard error of the mean), for the ‘intact’ and ‘impoverished’ condition in Experiment 1-3. Smaller dots denote individual participants’ means. Horizontal bars and * indicate significant differences between intact conditions and between impoverished conditions. **(B)** Distribution of *d’* for the yes-no animal detection task on the three impoverished sets of stimuli (phase-scramble, grayscale phase-scramble and texforms). Note that while phase-scramble and grayscale phase-scramble sets included 640 stimuli each, results for the texform set only consider the 350 “unrecognizable” stimuli that were used in Experiment 3. Box-plots represent the minimum, the first quartile, the median, the third quartile and the maximum of the population distribution. Dots represent each participant *d’* response. Cross represent the group average. Stars represent above-chance performance.

The effect of Image Type revealed that, in all experiments, the categorization-response was larger in the ‘intact’ than in the ‘impoverished’ condition. The effect of Experiment revealed that the categorization-response was larger in Experiment 1 than in Experiment 2 (*M_Difference_* ±*sd* = 0.10±0.08; 95% CI = 0.04 – 0.17; *t*(22) = 3.40; *P* = .003; *d* = 1.39) and in Experiment 1 than in Experiment 3 (*M_Difference_* ±*sd* = 0.07±0.05; 95% CI = 0.03 – 0.10; *t*(22) = 3.40; *P* = .003; *d* = 1.39), but did not differ between Experiments 2 and 3 (*M_Difference_*±*sd* = -0.04±0.07; 95% CI = -0.10 – 0.02; *t*(22) = -1.33; *P* = .20; *d* = 0.54).

Independent-sample *t*-tests further showed that for, the ‘intact’ sets, the response was greater in Experiments 1 v*s.* 2 (*M_Difference_*±*sd* = 0.12±0.13; 95% CI = 0.00 – 0.23; *t*(22) = 2.14; *P* = .044; *d* = 0.87), but did not differ between Experiments 1 and 3 (*M_Difference_*±*sd* = 0.05±0.08; 95% CI = -0.02 – 0.12; *t*(22) = 1.40; *P* = .18; *d* = 0.57), or between Experiments 2 and 3 (*M_Difference_*±*sd* = -0.07±0.14; 95% CI = -0.19 – 0.04; *t*(22) = -1.28; *P* = .21; *d* = 0.52). For the ‘impoverished’ sets, there was a significant difference between Experiments 1 and 2 (*M_Difference_*±*sd* = 0.09±0.06; 95% CI = 0.04 – 0.14; *t*(22) = 3.96; *P* < .001; *d* = 1.62) and Experiments 1 and 3 (*M_Difference_*±*sd* = 0.08±0.05; 95% CI = 0.04 – 0.13; *t*(22) = 3.88; *P* < .001; *d* = 1.58), but not between Experiments 2 and 3 (*M_Difference_*±*sd* = -0.01±0.05; 95% CI = -0.05 – 0.03; *t*(22) = -0.44; *P* = .66; *d* = 0.18).

### Experiment 4

Using a yes-no animal detection task, we tested whether low- and mid-level features preserved in unrecognizable images (and yielding an EEG categorization-response) supported accurate (above-chance; *P* < .017 corresponding to Bonferroni correction for 3 comparisons) animate-inanimate categorization, when an explicit judgment was required.

Results showed that categorization was at chance for grayscale phase-scramble stimuli (*M_d’_* = -0.01±0.02; 95% CI = -0.10 – Inf; *t*(19) = -0.30; *P* = 0.62; *d* = 0.07), but above chance for colorful phase-scramble stimuli (*M_d’_* = 0.12 ±0.20; 95% CI = 0.04 – Inf; *t*(19) = 2.74; *P* = 0.007; *d* = 0.61; Figure 3 B), and texforms (*M_d’_* = 0.74±0.26; 95% CI = 0.64 – Inf; *t*(19) = 12.69; *P* < 0.001; *d* = 2.84). Moreover, categorization was higher for texforms compared to phase-scramble stimuli (*M_difference_* = 0.62±0.23; 95% CI = 0.47 – 0.77; *t*(38) = 8.42; *P* < 0.001; *d* = 2.66) and to grayscale phase-scramble stimuli (*M_difference_* = 0.76±0.24; 95% CI = 0.61 – 0.91; *t*(38) = 10.07; *P* < 0.001; *d* = 3.18), but was not different for phase-scramble stimuli compare to grayscale phase-scramble stimuli (*M_difference_* = 0.14±0.21; 95% CI = 0.01 – 0.27; *t*(38) = 2.10; *P* = 0.042; *d* = 0.66).

The above analysis considered performance on all 640 stimuli for each set. Since in Experiment 3 we only included 350 images that turned out to be unrecognizable (i.e., named at the basic-level by less than 2 participants of an independent group of 19 participants), we repeated the analyses considering only this subset of texforms. Results remained unchanged. Categorization for texforms remained above chance (*M_d’_* = 0.53±0.22; 95% CI = 0.44 – Inf; *t*(19) = 10.97; *P* < 0.001; *d* = 2.45; Figure 3 B), and higher relative to phase-scramble (*M_difference_* = 0.44±0.21; 95% CI = 0.31 – 0.58; *t*(38) = 6.61; *P* < 0.001; *d* = 2.09) and grayscale phase-scramble stimuli (*M_difference_* = 0.55±0.22; 95% CI = 0.41 – 0.69; *t*(38) = 8.06; *P* < 0.001; *d* = 2.55). The distribution of classification accuracy values per stimulus are shown in Supplementary Figure 1.

In sum, participants’ performance in the yes-no animal detection task showed that color, contrast, luminance, texture, and global form (i.e., the features preserved in color phase-scramble images and texforms) all carry information that supports animate-inanimate classification. Among these, texture and global form preserved in texforms, appeared to provide the most reliable cues for explicit classification (see also Long et al., 2017), despite eliciting neural categorization responses that were markedly reduced relative to their intact counterparts. By contrast, color information preserved in color phase-scramble images, evoked the largest neural categorization response among the impoverished stimulus sets, although behavioral discrimination remained only slightly above the category boundary (*d′* = 0). Of the impoverished sets, grayscale phase-scramble images elicit the weakest neural categorization responses (approaching chance) and were not behaviorally discriminable. Overall, although both behavioral performance and neural categorization remained above chance for colored phase-scramble and texform stimuli, visual impoverishment substantially weakened both effects.

### Experiment 5

EEG results showed that categorization by animacy (i.e., the animate-inanimate categorization) emerges at different levels of the visual hierarchy (i.e., in higher-level anterior areas and early or middle areas of the ventral stream) as highlighted by the categorization effects obtained for both ‘intact’ and ‘impoverished’ stimuli. To probe this, we investigated which visual features, carried by different stimulus sets, distinguished between animate and inanimate objects, across different layers of an artificial DNN –VGG-19. In particular, we asked whether categorical information could be extracted in deeper as well as middle and first layers of the model from both ‘intact’ and ‘impoverished’ stimuli.

First, we validated our reference matrix (the representational dissimilarity matrix extracted from the output ‘fc8’ layer of VGG-19 for the original stimulus set, representing the network’s final object representation; Figure 4B, red box) by showing that it correlated strongly with a synthetic predictor matrix encoding the animate–inanimate distinction (ρ = 0.83, P < .001). Second, we found that the dissimilarity matrices for the other stimulus sets increasingly matched the reference matrix as stimulus complexity increased. In particular, correlations were the highest for the grayscale set (maximal correlation in layer fc8, *ρ* = .82, *P* < .001), although they were also significant for all other –more impoverished– sets (maximal correlation: phase-scramble, layer conv2_1, *ρ* = .34, *P* < .001; grayscale phase-scramble, layer conv2_2, *ρ* = .27, *P* < .001; texform, layer conv2_2, *ρ* = .26, *P* < .001) (see Supplementary Table S1).

**Figure 4.**
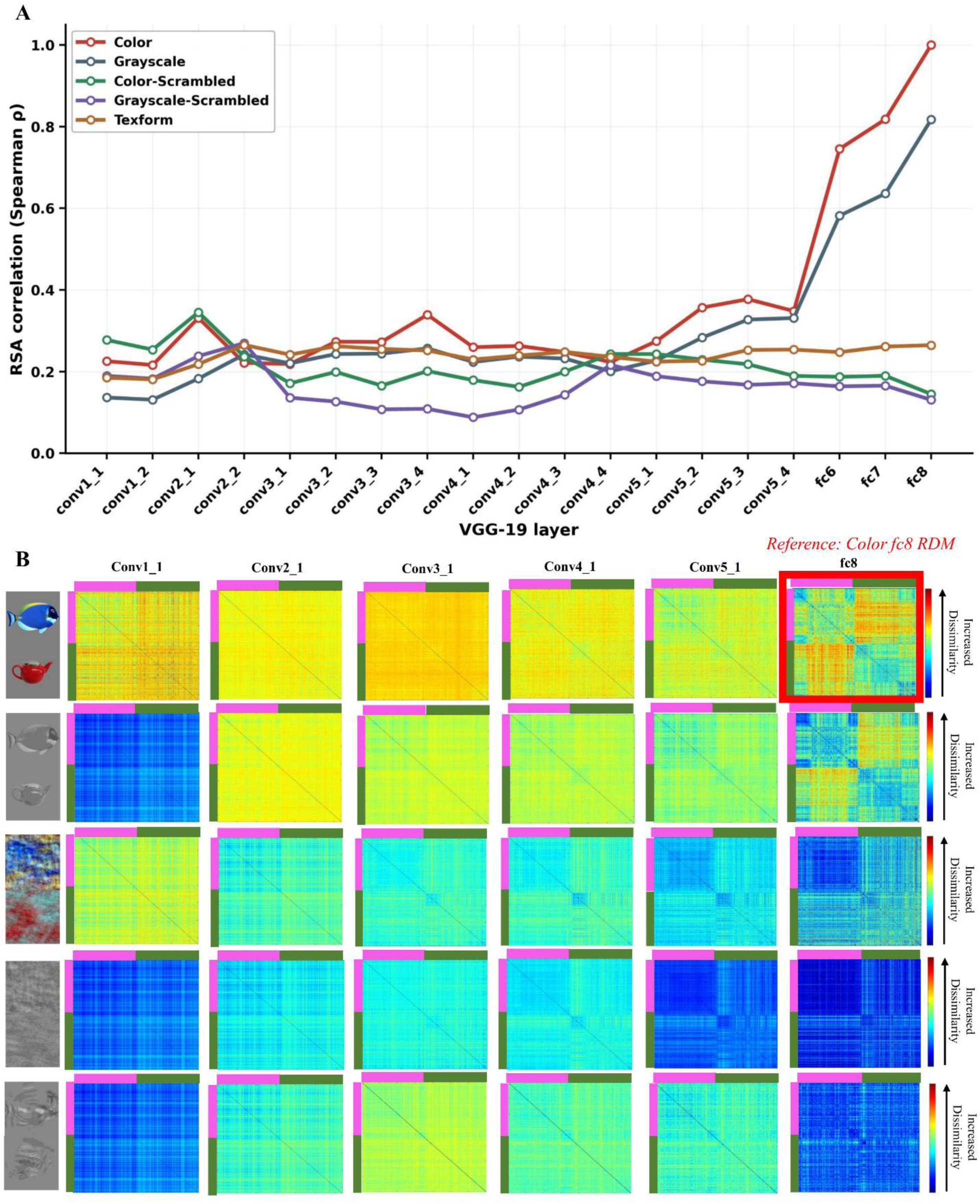
Correlation coefficients and dissimilarity matrices across image types and layers of VGG-19. **(A)**Layer-wise Spearman correlations between the RDM extracted for each stimulus set, from each VGG-19 layer and the reference RDM extracted from the fc8 layer for the intact color images (highlighted in red in B). Curves correspond to the intact color and grayscale set, color and grayscale phase-scramble sets and texform set. **(B)** Examples of RDM extracted from different layers for different sets. Rows correspond to different sets, columns correspond to different layers of VGG-19. Dissimilarity values range from 0 (lowest dissimilarity, dark blue) to 1 (highest dissimilarity, dark red). This figure illustrates the extent to which dissimilarity changed across layers and sets. The fc8 RDM for the intact color images corresponds to the reference matrix used for the correlations shown in A (see also Supplementary Table S1).

Third, we asked *where* accurate object representation emerged and peaked for the different stimulus sets (see Supplementary Table S1). For the intact sets (colorful and grayscale), coefficients of the correlation with the reference matrix increased as layers got deeper in the DNN, peaking in the last fully connected layers, with a significant difference across all layers of the two sets showing strongest correlations across layers for the intact colorful original stimulus set (*M_difference_* = 0.07 ±0.06; 95% CI = 0.07 – 0.10; *t*(18) = 4.64; *P* < 0.001; *d* = 1.07). For the impoverished sets, correlations with the reference matrix across layers were weaker, compared to the intact colorful set (phase-scramble: *M_difference_* = 0.16 ±0.25; 95% CI = 0.04 – 0.28; *t*(18) = 2.76; *P* = 0.013; *d* = .63; grayscale phase-scramble: *M_difference_* = 0.21 ±0.24; 95% CI = 0.09 – 0.32; *t*(18) = 3.81; *P* = 0.001; *d* = .88; texform: *M_difference_* = 0.13 ±0.22; 95% CI = 0.03 – 0.24; *t*(18) = 2.61; *P* = 0.018; *d* = .60). For the phase-scramble and texform sets, correlation coefficients did not change much from the first and middle layers to the deeper layers (Figure 4A), suggesting that the information extracted in deeper layers did not add to the representation of those images. In some cases (i.e., for both phase-scramble sets), correlation coefficients were even higher in middle than deepest layers, suggesting that middle layers were better tuned to the features carried by those stimuli. These results show that object representation embedding the animate-inanimate distinction emerges across different processing stages of the visual hierarchy, relying on high-level, as well as mid- and low-level visual features. These results were qualitatively replicated across additional DNN architectures such as AlexNet, ResNet-50, CORnet-S, and VGG-16 (see Supplementary Materials and Supplementary Table S2).

## Discussion

The animate-inanimate distinction underlies *life detection*, a capacity that is crucial for the survival of animal species. A rich body of work has shown that, in visual cortex and visual perception, objects are represented as (more or less) animate, in line with intuitive taxonomies, by virtue of complex properties, such as the presence of a face or body, being alive, possessing agency and/or being unpredictable, among others (Bracci & Op De Beeck, 2023; Jozwik et al., 2022; Leys et al., 2025; Papeo et al., 2017; Ritchie et al., 2021; Thorat et al., 2019). Another line of research has shown that the animate-inanimate distinction can already emerge from mid-level features such as shape and texture, and low-level feature such as color and power-spectrum information (Ayzenberg & Behrmann, 2022; Coggan et al., 2016; Grootswagers et al., 2017; Loke et al., 2025; Long et al., 2017, 2018; Peykarjou et al., 2024a; Wang et al., 2022).

Building on this work, we used ftEEG to investigate whether the neural response time-locked to stimulus presentation already encodes the animate-inanimate distinction. We further quantified the extent to which mid- and low-level visual features support this distinction in neural activity and explicit behavioral judgements, when they constituted the only diagnostic information available. To this end, we used stimuli that preserved selected (low- or mid-level) visual properties while remaining unrecognizable at the basic level, thereby dissociating visual information supporting the animate–inanimate distinction from explicit object recognition. We sampled the diversity of real-world objects using an unprecedentedly large and heterogeneous stimulus set (320 animals and 320 inanimate objects) including mammals, birds, fish and amphibians, furniture, vehicles, tools, plants and vegetables. This design increased the likelihood that the observed categorization response reflected category-level visual information rather than idiosyncratic properties of particular stimuli.

ftEEG results demonstrated that information relevant to the animate–inanimate distinction is extracted rapidly during object perception, as reflected by neural responses time-locked to stimulus presentation. This process appears to be largely automatic: participants received no instructions to categorize the stimuli, were not informed about the periodic categorical changes in the stimulus sequences, and variations in attention—as indexed by post-sequence task performance and base-frequency response amplitudes (see Results and Supplementary Materials, pp. 1–3)—did not account for differences in the categorization response. Although participants’ awareness of the categorical manipulation was not formally assessed, these findings suggest that the animate– inanimate distinction emerges without explicit categorization demands.

Moreover, ftEEG results demonstrated that information relevant for the animate-inanimate distinction can be extracted from both mid- and low-level visual features. In particular, a strong, widespread response to periodic categorical changes was elicited by colorful, intact, images of real-world objects (Experiment 1). This response was only moderately reduced when color, luminance and contrast information was removed (i.e., grayscale intact images; Experiment 2), and remained detectable, although markedly weaker, for unrecognizable images preserving only low-level spectral information (color, contrast, luminance, number of pixels, power spectrum, in the phase-scramble set) or mid-level information (texture and global form in texforms) (Experiments 1-3). Above-chance neural discrimination for the impoverished sets (color phase-scramble and texform sets) was accompanied by above-chance behavioral classification (Experiment 4), suggesting that these neural representations contained sufficient information to support explicit animate-inanimate judgements. By contrast, the weak, near-chance, neural response elicited by grayscale phase-scramble stimuli did not appear to provide sufficient information for explicit classification, which remained at chance.

In sum, combined EEG and behavioral data show that visual impoverishment substantially attenuates both neural and behavioral animate-inanimate categorization. Nevertheless, above-chance effects for phase-scramble and texform stimuli demonstrate that color, contrast, luminance, texture, and global form contribute information that supports animate–inanimate classification during visual processing and in explicit behavioral judgments.

The fact that phase-scramble and texform stimuli were unrecognizable at the basic level indicates that the animate-inanimate distinction did not depend on explicit object recognition and is unlikely to have arisen from top-down feedback following object identification. Instead, our findings suggest that category-relevant information is extracted progressively across successive stages of visual processing, with the accumulation of visual information from low- to high-level features strengthening the animate–inanimate distinction, as reflected in the increasing amplitude of the EEG categorization response. Indeed, our approach, in which the amplitude of the EEG categorization-response is taken as an index of category discriminability, suggests a hierarchy of visual distinctiveness, whereby higher-level features (in the ‘intact’ sets) provide stronger categorical information than the isolated mid- and low-level features preserved in the impoverished sets. Moreover, although, mid-level features (texforms) did not elicit significantly larger neural categorization responses than low-level features (in phase-scramble sets) (Figure 3 A), they proved more informative for explicit judgments. Among the low- and mid-level features, color provided a substantial additional benefit: preserving color increased the categorization response in both intact and phase-scramble stimuli and enabled above-chance performance in the yes–no animal detection task with phase-scramble images.

EEG and behavioral findings were complemented by analyses of VGG-19 and other DNNs. In artificial networks, as in human vision, representations matching the network’s final object representation emerged already in early and intermediate layers. This supports a model where visual information preserved in the impoverished images is sufficient to establish an animate– inanimate representational structure across successive stages of hierarchical processing. Moreover, our findings suggest a convergence between biological and artificial vision in the representation of the animate–inanimate distinction, a relationship that has remained ambiguous in previous studies (Bracci et al., 2019; Duyck et al., 2024).

Our work raises an important question about the level of categorization supported by low- and mid-level visual features. Our results show that these features are sufficient to support to superordinate (animate-inanimate) categorization, even when they do not support basic-level object recognition. In the preliminary validation of our stimuli, participants were asked to name texforms and phase-scramble images. Although visual typically favors the basic level (e.g., “zucchini” rather than “vegetable” or “inanimate object”; Mandelbaum, 2018; Murphy & Brownell, 1985; Rogers & Patterson, 2007; Rosch, 1978), none of the phase-scramble images and only a small subset of texforms (excluded from Experiment 3) were recognized at the basic level. Nevertheless, performance with the same stimuli in the yes-or-no animal detection tasks was significantly above-chance, meaning that superordinate animate-inanimate categorization was possible, where basic-level recognition was not.

Naming and yes-or-no discrimination are different tasks. Therefore, it remains possible that participants could discriminate between basic-level categories under forced-choice conditions (e.g., cat versus dog). However, it also remains possible that impoverished visual information provides preferential access to superordinate category representations. This interpretation is consistent with developmental results showing that, while young (4-month-old) infants can represent the broad animate-inanimate distinction, basic-level categorical distinctions develops later and appear to require the integration of richer visual features across the visual cortex (Spriet et al., 2022). Determining how the complexity of visual features, their cortical representations, and the level of categorization (superordinate versus basic) interact remains an important avenue for future research.

Finally, our study did not address the representational organization along the *animacy continuum*, as proposed by a growing body of work (Bracci & Op De Beeck, 2023; Connolly et al., 2012; Jozwik et al., 2022; Leys et al., 2025; Papeo et al., 2017; Ritchie et al., 2021; Sha et al., 2015; Thorat et al., 2019). A more comprehensive understanding of visual object categorization will require determining whether, and how, the basic visual features investigated here relate to or interact with the more complex properties proposed to organize this continuum (e.g., presence of a face or body, being alive, agency, or unpredictability). Addressing this question remains an important direction for future research.

In conclusion, we showed that the animate-inanimate distinction is pervasive in the processing of visual stimuli, and is remarkably resilient to the loss of information. Low- and mid-level features, encoded in early and intermediate aspects of the visual ventral stream for object recognition, are sufficient to elicit rapid, automatic neural categorization response, and to support explicit animate-inanimate judgments. However, impoverished stimuli evoked substantially weaker categorization response than intact objects, and could not be recognized beyond a coarse animate-inanimate distinction. These findings support a view of visual categorization as an incremental process in which increasingly complex visual features are progressively integrated across the visual cortex to strengthen categorical representations. On this view, broad distinctions such as animate *vs.* inanimate, which can rely on relatively simple visual features, may emerge before finer-grained object categories become available during visual processing.

## Data availability

Stimuli, EEG data and code for the main analyses are available here: https://osf.io/e6fjn/overview?view_only=f64fc9e1f3ce4d49b3aec428beeea238.

## Authors’ contributions - CRediT

*Céline Spriet*: Conceptualization, Methodology, Formal Analysis, Investigation, Data Curation, Writing – Original Draft, Writing – Review & Editing, Visualization; *Farzad Rostami*: Formal Analysis, Data Curation, Writing – Original Draft, Writing – Review & Editing, Visualization; *Jean-Rémy Hochmann*: Conceptualization, Methodology, Writing – Review & Editing, Supervision; *Liuba Papeo*: Conceptualization, Methodology, Writing – Review & Editing, Supervision, Funding Acquisition.

## Acknowledgments

We thank Sofie Vettori for helping with data collection. L. P. was supported by an ANR-DFG Franco-German Grant (FRAL_RELATIONS_268980). C.S. was supported by a fellowship of the Fondation pour la Recherche Médicale (FDT202304016547). F.R. was supported by a fellowship of the LabEx CORTEX of the University of Lyon (ANR-11-LABX-0042).

## Citation diversity statement

The proportion of authorship teams in this paper is M/M = .357, W/M = .321, M/W = .196, and W/W = .125.

