## Supplementary Material for "Animate-inanimate object categorization from minimal visual information in the human brain, human behavior, and deep neural networks"

Céline Spriet[1], Farzad Rostami[1], Jean-Rémy Hochmann[1], Liuba Papeo[1]

[1]*Institut des Sciences Cognitives—Marc Jeannerod, UMR5229, Centre National de la Recherche Scientifique (CNRS) and Université Claude Bernard Lyon 1, 67 Bd. Pinel – 69675 Bron (France)*

Correspondence to:

C.S.

L.P.

**Supplementary material**

- Supplementary results
- Supplementary figure 1
- Supplementary table S1
- Supplementary table S2
- Supplementary table S3

### 1. Post-sequence task

To motivate participants to pay attention to the stimuli presented in rapid sequence during ftEEG, at the end of each ftEEG trial, participants were tasked with a recognition test, in which they had to decide whether a target-image was shown of the last trial. We examined the participants' performance (accuracy rates) using a one-sided *t*-test against chance-level (0.5%) and correcting for multiple comparisons using the Bonferroni correction ($P < .008$ corresponding to 6 performed tests).

Accuracy was above chance only for the intact condition (original stimulus set) of Experiment 1 ($M_{Accuracy} \pm sd$ = 0.56±0.07; 95% CI = 0.52 – Inf; $t(11) = 2.88$; $P = .008$; $d = 0.83$) , whereas it remained at chance level in all other conditions and experiments (Experiment 1, impoverished: $M_{Accuracy} \pm sd$ = 0.54±0.07; 95% CI = 0.51 – Inf; $t(11) = 2.05$; $P = .03$; $d = 0.59$; Experiment 2, intact: $M_{Accuracy} \pm sd$ = 0.55±0.10; 95% CI = 0.50 – Inf; $t(11) = 1.79$; $P = .050$; $d$ = 0.52; impoverished: $M_{Accuracy} \pm sd$ = 0.51±0.09; 95% CI = 0.46 – Inf; $t(11) = 0.29$; $P = .39$; $d = 0.08$; Experiment 3, intact: $M_{Accuracy} \pm sd$ = 0.49±0.09; 95% CI = 0.45 – Inf; $t(11) = -0.21$; $P = .58$; $d$ = 0.60; impoverished: $M_{Accuracy} \pm sd$ = 0.47±0.09; 95% CI = 0.42 – Inf; $t(11) = -1.16$; $P = .86$; $d$ = 0.34).

In sum, the intact condition in Experiment 1 was the only condition in which performance on the post-sequence task exceeded chance. However, significant neural categorization responses—including responses comparable in magnitude to those observed in the intact condition of Experiment 1 (e.g., the intact condition of Experiment 3)—were also obtained when post-sequence performance remained at chance. Thus, even if some stimulus sets (e.g., the intact images in Experiment 1) attracted greater attention or were easier to process and remember, these differences could not account for the pattern of oddball categorization responses observed across Experiments 1–3.

### 2. Comparisons of the base-rate responses

We took the base-frequency responses as a possible index of attention to the visual stimulation and examined whether differences between conditions in the base-frequency responses could reflect the differences in the oddball responses, thus suggesting a link between attention to the stimuli and categorization. For each experiment, base-rate responses (baseline-subtracted amplitudes summed over the 8 harmonics and averaged over all 128 electrodes) were compared between conditions using *t* tests.

Base-frequency responses did not differ between intact and impoverished conditions in Experiment 1 ($M_{Difference} \pm SD = 0.03 \pm 0.25$; 95% CI = −0.13 to 0.19; $t(11) = 0.46$; $P = .66$; $d = 0.13$) or Experiment 2 ($M_{Difference} \pm SD = 0.09 \pm 0.14$; 95% CI = −0.00 to 0.18; $t(11) = 2.20$; $P = .05$; $d = 0.63$), and differed only in Experiment 3, where intact grayscale images elicited larger responses than texforms ($M_{Difference} \pm SD = 0.15 \pm 0.17$; 95% CI = 0.04 to 0.26; $t(11) = 3.10$; $P = .01$; $d = 0.90$). Importantly, categorization responses were reduced for the impoverished stimuli in all three experiments, including Experiments 1 and 2, where base-frequency responses were unchanged. Thus, even if some stimulus sets (e.g., intact images) elicited greater attention, as indexed by larger base-frequency responses, differences in the base response to visual stimulation cannot account for the pattern of oddball categorization responses observed in Experiments 1-3.

### 3. Accuracy per stimulus in the yes-no animal detection task

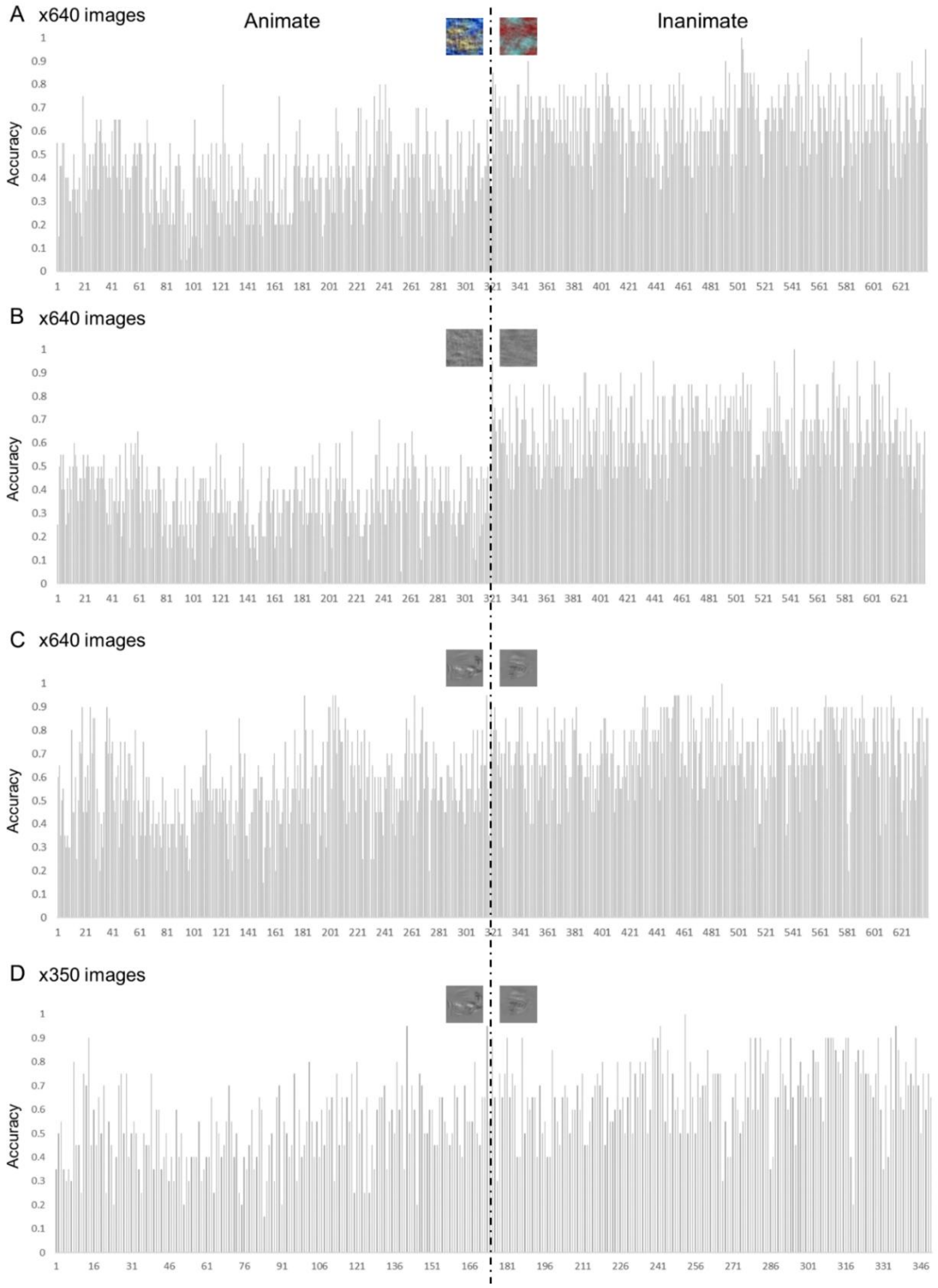


**Figure S1. Accuracy per stimulus.** By-stimulus accuracy for the 640 phase-scramble stimuli **(A)**, 640 grayscale phase-scramble stimuli **(B)**, 640 texform stimuli **(C)** and the subset of 350 texform stimuli used in Experiment 3 **(D)**. Overall, inanimate objects were more frequently classified correctly relative to animate objects across all stimulus sets.

### 4. Multi-architecture RSA

We measured the correlation between the reference-matrix (i.e., the representational dissimilarity matrix extracted from the output 'fc8' layer of VGG-19 for the original stimulus set, representing the network's final object representation) and the matrices representing stimulus representation at each layer of VGG-19, for the different stimulus sets (Table S1). We found significant correlations for all layers and all stimulus sets, the highest correlation being with the matrices obtained for the higher-level layers of the model, with the grayscale stimuli.

**Table S1.** Correlations between the *reference-matrix* extracted for the intact colorful original set, from the last layer fc8 (highlighted with the frame), and the matrix extracted for each stimulus set, from each layer of VGG-19.

| | Original intact | | Grayscale intact | | Phase-scramble (PS) | | Grayscale PS | | Texform | |
|---|---|---|---|---|---|---|---|---|---|---|
| Layers | *ρ* | *P* | *ρ* | *P* | *ρ* | *P* | *ρ* | *P* | *ρ* | *P* |
| conv1_1 | .225 | **<.001** | .136 | **<.001** | .277 | **<.001** | .190 | **<.001** | .185 | **<.001** |
| conv1_2 | .216 | **<.001** | .131 | **<.001** | .254 | **<.001** | .183 | **<.001** | .181 | **<.001** |
| conv2_1 | .331 | **<.001** | .183 | **<.001** | .345 | **<.001** | .238 | **<.001** | .218 | **<.001** |
| conv2_2 | .221 | **<.001** | .242 | **<.001** | .237 | **<.001** | .270 | **<.001** | .265 | **<.001** |
| conv3_1 | .218 | **<.001** | .220 | **<.001** | .171 | **<.001** | .136 | **<.001** | .241 | **<.001** |
| conv3_2 | .273 | **<.001** | .243 | **<.001** | .199 | **<.001** | .126 | **<.001** | .262 | **<.001** |
| conv3_3 | .273 | **<.001** | .244 | **<.001** | .166 | **<.001** | .107 | **<.001** | .255 | **<.001** |
| conv3_4 | .339 | **<.001** | .257 | **<.001** | .201 | **<.001** | .109 | **<.001** | .252 | **<.001** |
| conv4_1 | .260 | **<.001** | .224 | **<.001** | .179 | **<.001** | .088 | **<.001** | .229 | **<.001** |
| conv4_2 | .263 | **<.001** | .237 | **<.001** | .162 | **<.001** | .107 | **<.001** | .239 | **<.001** |
| conv4_3 | .248 | **<.001** | .232 | **<.001** | .200 | **<.001** | .143 | **<.001** | .248 | **<.001** |
| conv4_4 | .223 | **<.001** | .201 | **<.001** | .243 | **<.001** | .216 | **<.001** | .236 | **<.001** |
| conv5_1 | .274 | **<.001** | .227 | **<.001** | .243 | **<.001** | .189 | **<.001** | .224 | **<.001** |
| conv5_2 | .357 | **<.001** | .283 | **<.001** | .229 | **<.001** | .176 | **<.001** | .227 | **<.001** |
| conv5_3 | .377 | **<.001** | .327 | **<.001** | .218 | **<.001** | .168 | **<.001** | .253 | **<.001** |
| conv5_4 | .348 | **<.001** | .331 | **<.001** | .189 | **<.001** | .171 | **<.001** | .254 | **<.001** |
| fc6 | .745 | **<.001** | .582 | **<.001** | .187 | **<.001** | .164 | **<.001** | .248 | **<.001** |
| fc7 | .818 | **<.001** | .636 | **<.001** | .190 | **<.001** | .166 | **<.001** | .261 | **<.001** |
| fc8 | 1.000 | **<.001** | .817 | **<.001** | .145 | **<.001** | .131 | **<.001** | .265 | **<.001** |

*Note: Colors denote the strength of the correlation from very strong (red: ρ = 0.8 to 1), to increasingly weaker (orange: ρ = 0.6 to 0.79; yellow, ρ = 0.4 to 0.59; green, ρ = 0.2 to 0.39; blue, ρ = 0 to 0.19). Statistical significance was assessed using 1000 row/column permutations of the reference RDM, and P-values were corrected for multiple comparisons across the layer × stimulus-set RSA correlations using the Benjamini–Hochberg false discovery rate procedure. PS: phase-scramble.*

We further assessed whether the DNN representations contained explicit animate–inanimate categorical structure by correlating each VGG-19-layer RDM with a theoretical binary animate–inanimate model RDM (Table S2). In this binary model, cells representing comparisons within-category were assigned 0 and cells representing comparisons between-category were assigned 1. For each stimulus set and each VGG-19-layer, Spearman correlations were computed over the upper triangular RDM entries, excluding the diagonal. Statistical significance was assessed using 1,000 label permutations preserving the animate/inanimate category counts, followed by Benjamini–Hochberg FDR correction across the layer × stimulus-set RSA correlations. Results showed that the Original intact and Grayscale intact stimulus sets carried the strongest animate–inanimate categorical structure in deeper layers of VGG-19. Phase-scramble, grayscale phase-scramble, and texform stimuli showed weaker correlations, indicating that visual impoverishment weakened the animate–inanimate representational structure in the network.

**Table S2.** Correlations between a binary animate–inanimate model RDM and the matrix extracted from each stimulus set, from each layer of VGG-19. In the binary model RDM, within-category image pairs were assigned a dissimilarity of 0 and between-category animate–inanimate pairs were assigned a dissimilarity of 1.

| | Original intact | | Grayscale intact | | Phase-scramble (PS) | | Grayscale PS | | Texform | |
|---|---|---|---|---|---|---|---|---|---|---|
| Layers | *ρ* | *P* | *ρ* | *P* | *ρ* | *P* | *ρ* | *P* | *ρ* | *P* |
| conv1_1 | .050 | **<.001** | .017 | **<.001** | .045 | **<.001** | .017 | **<.001** | .016 | **<.001** |
| conv1_2 | .041 | **<.001** | .020 | **<.001** | .034 | **<.001** | .020 | **<.001** | .019 | **<.001** |
| conv2_1 | .029 | **<.001** | .022 | **<.001** | .036 | **<.001** | .021 | **<.001** | .032 | **<.001** |
| conv2_2 | .026 | **<.001** | .021 | **<.001** | .017 | **<.001** | .021 | **<.001** | .031 | **<.001** |
| conv3_1 | .053 | **<.001** | .050 | **<.001** | .058 | **<.001** | .056 | **<.001** | .070 | **<.001** |
| conv3_2 | .114 | **<.001** | .056 | **<.001** | .090 | **<.001** | .057 | **<.001** | .095 | **<.001** |
| conv3_3 | .122 | **<.001** | .050 | **<.001** | .078 | **<.001** | .056 | **<.001** | .104 | **<.001** |
| conv3_4 | .137 | **<.001** | .045 | **<.001** | .098 | **<.001** | .055 | **<.001** | .112 | **<.001** |
| conv4_1 | 168 | **<.001** | .082 | **<.001** | .109 | **<.001** | .078 | **<.001** | .132 | **<.001** |
| conv4_2 | .166 | **<.001** | .069 | **<.001** | .097 | **<.001** | .062 | **<.001** | .140 | **<.001** |
| conv4_3 | .189 | **<.001** | .068 | **<.001** | .085 | **<.001** | .058 | **<.001** | .156 | **<.001** |
| conv4_4 | .165 | **<.001** | .054 | **<.001** | .061 | **<.001** | .045 | **<.001** | .157 | **<.001** |
| conv5_1 | .204 | **<.001** | .107 | **<.001** | .128 | **<.001** | .067 | **<.001** | .192 | **<.001** |
| conv5_2 | .211 | **<.001** | .151 | **<.001** | .183 | **<.001** | .063 | **<.001** | .189 | **<.001** |
| conv5_3 | .191 | **<.001** | .177 | **<.001** | .199 | **<.001** | .074 | **<.001** | .169 | **<.001** |
| conv5_4 | .185 | **<.001** | .198 | **<.001** | .176 | **<.001** | .073 | **<.001** | .171 | **<.001** |
| fc6 | .362 | **<.001** | .325 | **<.001** | .211 | **<.001** | .103 | **<.001** | .153 | **<.001** |
| fc7 | .265 | **<.001** | .276 | **<.001** | .203 | **<.001** | .111 | **<.001** | .148 | **<.001** |
| fc8 | .508 | **<.001** | .427 | **<.001** | .172 | **<.001** | .123 | **<.001** | .121 | **<.001** |

*Note: Colors denote the strength of the correlation based on a consistent scale, from very strong (red: ρ = 0.8 to 1; orange: ρ = 0.6 to 0.79; yellow: ρ = 0.4 to 0.59), to increasingly weaker (green: ρ = 0.2 to 0.39; blue: ρ = 0.1 to 0.19), and very weak (light purple: ρ = 0 to 0.09). Statistical significance was assessed using 1000 row/column permutations of the reference RDM, and P-values were corrected for multiple comparisons across the layer × stimulus-set RSA correlations using the Benjamini–Hochberg false discovery rate procedure. PS: phase-scramble.*

In addition to the VGG-19 analysis, we performed a supplementary multi-architecture RSA analysis using AlexNet, ResNet-50, CORnet-S, and VGG-16 and the THINGvision package in python (https://thingsvision.github.io/). For these analyses, we relied on correlations with a reference RDM analysis (rather than a binary animate-inanimate categorical model) as we considered that the reference RDM provides a richer benchmark preserving graded similarity relations among individual exemplars, in addition to the broad animate–inanimate separation. For each architecture, we extracted layer-wise activations for the same five stimulus sets used in the original DNN analysis: original color images, grayscale images, phase-scramble images, grayscale phase-scramble images, and texforms. For each model, layer, and stimulus set, we constructed a representational dissimilarity matrix (RDM) using 1 − Pearson correlation between activation vectors. We then correlated each layer/stimulus-set RDM with the final-layer RDM of the corresponding model, reflecting the representation of original intact stimulus set, using Spearman correlations.

Image order was deterministic across architectures, with animate images followed by inanimate images and numeric filename order within category. Significance was assessed using 1000 row/column permutations of the reference RDM, followed by Benjamini–Hochberg FDR correction within model. For each model and stimulus set, we identified the layer showing the strongest correlation with that model's reference RDM, i.e., the last layer's RDM obtained from the original colorful stimulus set (see Supplementary Table S3). Notably, while CORnet-S yielded the strongest representational alignment for intact grayscale images, AlexNet showed the highest correlations across all three impoverished stimulus sets (phase-scramble, grayscale phase-scramble, and texforms). Overall, the results reproduced the qualitative pattern observed with VGG-19: grayscale images remained strongly aligned with the original representation, whereas phase-scramble, grayscale phase-scramble, and texform images showed weaker representational alignment. These analyses show that the qualitative pattern observed with the VGG-19 generalizes across different architectures.

**Table S3.** Supplementary multi-architecture within-model RSA. For each stimulus set, the values indicate the exact maximal representational alignment (Spearman ρ) and the corresponding best layer for AlexNet, ResNet-50, CORnet-S, and VGG-16. For each architecture, the reference was the model's final-layer RDM obtained with the original stimulus set.

| **Stimulus set** | **Best-layer Spearman ρ** | **Best layer by model** |
|---|---|---|
| **Original (Exp. 1)** | AlexNet: 1.000*<br>CORnet-S: 1.000*<br>ResNet-50: 1.000*<br>VGG-16: 1.000* | AlexNet: classifier.6<br>CORnet-S: decoder.linear<br>ResNet-50: fc<br>VGG-16: classifier.6 |
| **Grayscale (Exps. 2-3)** | AlexNet: 0.735<br>CORnet-S: 0.838<br>ResNet-50: 0.710<br>VGG-16: 0.805 | AlexNet: classifier.6<br>CORnet-S: decoder.linear<br>ResNet-50: fc<br>VGG-16: classifier.6 |
| **Phase-scramble (Exp. 1)** | AlexNet: 0.326<br>CORnet-S: 0.253<br>ResNet-50: 0.211<br>VGG-16: 0.228 | AlexNet: features.11<br>CORnet-S: decoder.avgpool<br>ResNet-50: layer2<br>VGG-16: features.30 |
| **Grayscale phase-scramble (Exp. 2)** | AlexNet: 0.257<br>CORnet-S: 0.193<br>ResNet-50: 0.230<br>VGG-16: 0.164 | AlexNet: features.9<br>CORnet-S: decoder.linear<br>ResNet-50: layer2<br>VGG-16: features.30 |
| **Texform (Exp. 3)** | AlexNet: 0.341<br>CORnet-S: 0.177<br>ResNet-50: 0.212<br>VGG-16: 0.160 | AlexNet: classifier.6<br>CORnet-S: decoder.linear<br>ResNet-50: layer2<br>VGG-16: classifier.6 |

** Original stimulus set equals 1.000 by construction because each model's final-layer original RDM served as the reference model. For the non-reference stimulus sets, all reported best-layer correlations survived FDR correction, q < .001; this threshold corresponds to the permutation floor for 1000 permutations.*
